# A Novel Machine Learning Strategy for Prediction of Antihypertensive Peptides Derived from Food with High Efficiency

**DOI:** 10.1101/2020.08.12.248955

**Authors:** Liyang Wang, Dantong Niu, Xiaoya Wang, Qun Shen, Yong Xue

**Affiliations:** College of Food Science and Nutritional Engineering, China Agricultural University, Beijing 100083, China; (L.W.); (X.W.); (Q.S.); (Y.X.); EECS Department, University of California, Berkeley, Berkeley, CA 94720-1770; (D.N.)

**Keywords:** ACE inhibitory peptides, XGBoost model, peptide-protein technology, protein from bovine milk, high-speed screening

## Abstract

Strategies to screen antihypertensive peptides with high throughput and rapid speed will be doubtlessly contributed to the treatment of hypertension. The food-derived antihypertensive peptides can reduce blood pressure without side effects. In present study, a novel model based on Extreme Gradient Boosting (XGBoost) algorithm was developed using the primary structural features of the food-derived peptides, and its performance in the prediction of antihypertensive peptides was compared with the dominating machine learning models. To further reflect the reliability of the method in real situation, the optimized XGBoost model was utilized to predict the antihypertensive degree of k-mer peptides cutting from 6 key proteins in bovine milk and the peptide-protein docking technology was introduced to verify the findings. The results showed that the XGBoost model achieved outstanding performance with the accuracy of 0.9841 and the area under the receiver operating characteristic curve of 0.9428, which were better than the other models. Using the XGBoost model, the prediction of antihypertensive peptides derived from milk protein was consistent with the peptide-protein docking results, and was more efficient. Our results indicate that using XGBoost algorithm as a novel auxiliary tool is feasible for screening antihypertensive peptide derived from food with high throughput and high efficiency.

## 1 Introduction

Hypertension, also known as cardiovascular syndrome, is a modifiable risk factor responsible for a high burden of disability and death [1]. The heart and blood vessels are the main target organs, suffering from the pathologically and physiologically negative effects of hypertension. In the early stage, hypertension patients may behave no obvious symptoms, but long-term high blood pressure will cause ventricular hypertrophy and enlargement, and even burden the arteries, causing the stiffness of arterial wall and the reduction of arterial diameter, therefore endangering the blood supply of corresponding organs such as heart, brain and kidney [2–5]. In the final stage, considering that hypertension patients are always accompanied by other metabolic diseases such as diabetes or hypercholesterolemia, serious complications of the above-mentioned organs will occur, which pose threat to health of humans and aggravate the formation of atherosclerosis. Therefore, the effective treatment of hypertension should be paid more attention. Although drug intervention is the conventional treatment for hypertension, there is increasing evidence to indicate that non-pharmacologic strategies of anti-hypertension such as dietary modifications could have a considerable effect in the prevention and treatment of hypertension [6–8].

Currently, the common antihypertensive drugs for clinical treatment are diuretics, adrenergic receptor blockers, calcium channel blockers and angiotensin converting enzyme (ACE) inhibitors. Despite of satisfactory antihypertensive effect, they inevitably have more or less side effects or adverse reactions. On the contrary, food-derived antihypertensive peptides, a specific class of micro-molecule peptides, can reduce blood pressure of humans with many attractive advantages. For example, they have outstanding performance while cause no excessive reduction of blood pressure. Moreover, the properties of them have no toxicity, side effects and adverse reactions, which bring no harmful effect to people with normal and abnormal blood pressure. Therefore, the discovery and verification of these peptides have been brought to the hot spot of current research [9,10]. ACE, as a zinc-containing dipeptide carboxypeptidase, plays a vital role in the regulatory mechanism of blood pressure including renin angiotensin regulatory system (RAS) and kallikrein kinin system (KKS), so that inhibiting ACE activity is considered as a key measure to treat hypertension and ACE inhibitors become as the important targets when doing drug screening. Currently, a large number of highly effective and specific ACE inhibitors including some bioactive peptides have functioned in the clinical treatment of hypertension and congestive heart failure. It is worth noting that considerable amount of ACE inhibitory peptides have been discovered in food. These biologically active peptides compete and combine with Zn^2+^ at the active site of ACE to inhibit its activity and thereby suppress the conversion of angiotensin I. At the same time, such process induced by food-derived antihypertensive peptides is a natural process of inhibition with less toxic. The traditional screening methods for ACE inhibitory peptides mainly depend on *in vitro* or/and *in vivo* experiments [11], which are time-consuming and expensive. Thus, the design and establishment of a screening method with high throughput and speediness have extremely important for antihypertensive peptides.

With the development of artificial intelligence technology, machine learning has been applied in the field of drug screening, which could shorten the experiment time and make a great achievement to a large degree by studying the characteristics of known drug molecules. There have been many reports on the development and research of ACE inhibitory peptides [12,13]. These studies have achieved relatively good results mainly by employing quantitative structure activity relationship (QSAR) technique and traditional machine learning models. In the screening tasks of other targets [14–16], researchers also adopted machine learning strategies (including deep learning) to screen inhibitors of key targets, such as dipeptidyl peptidase-4 (DPP-IV) inhibitors and anticancer peptide (ACP) inhibitors. As for the study of peptide activity prediction, the feature extraction of the primary structure is very important. Dubey at al. [17] extracted the four proteins (MN, NO, MO, and MNO) in HIV-1 by using amino acid composition (AAC), and then the support vector machine (SVM) model was used to make classifications, which performing with high accuracy. Chou et al. [18,19] proposed a method named as PseAAC to represent the feature of amino acid composition, which combines its physical and chemical properties, and this method has been widely used in bioinformatics of protein function.

In this study, we proposed to use PseAAC to exploit the features extracted from ACE inhibitory peptides (positive samples) and the peptides (negative samples) randomly selected from the UniProt library, and then designed a parameter-optimized XGBoost machine learning model to train the data and perform predictions. Though the XGBoost has been widely used in many fields because of its superior structure [20,21], such utilizations have been rarely found in the research of drug screening. In our study, three public databases of ACE inhibitory peptides were explored and some dominating machine learning methods presented in previous researches were compared with our XGBoost model. Through 5-fold cross-validation, we found that the XGBoost model was demonstrated with superior performance in the screening task of ACE inhibitory peptides. In order to verify the generalization ability of this method in real situation, we initially selected 6 proteins with unambiguous structure in bovine milk as row sequences to get k-mer peptides using cycle cutting algorithm. After predicting by the XGBoost model, candidate inhibitory and non-inhibitory peptides were further confirmed by using the peptide-protein docking technology. The workflow of our study is shown in **Figure** 1. It is novel method to screen ACE inhibitory peptides by using XGBoost algorithm and PseAAC based on the feature of amino acid composition. And it is the first time to completely screen antihypertensive peptides based on food protein with unambiguous structure, which exhibited with high throughput and high efficiency. Furthermore, the method of XGBoost algorithm performed better than other machine learning algorithms, which would facilitate the rapid screen of food-derived ACE inhibitory peptides in future.

**Figure 1:**
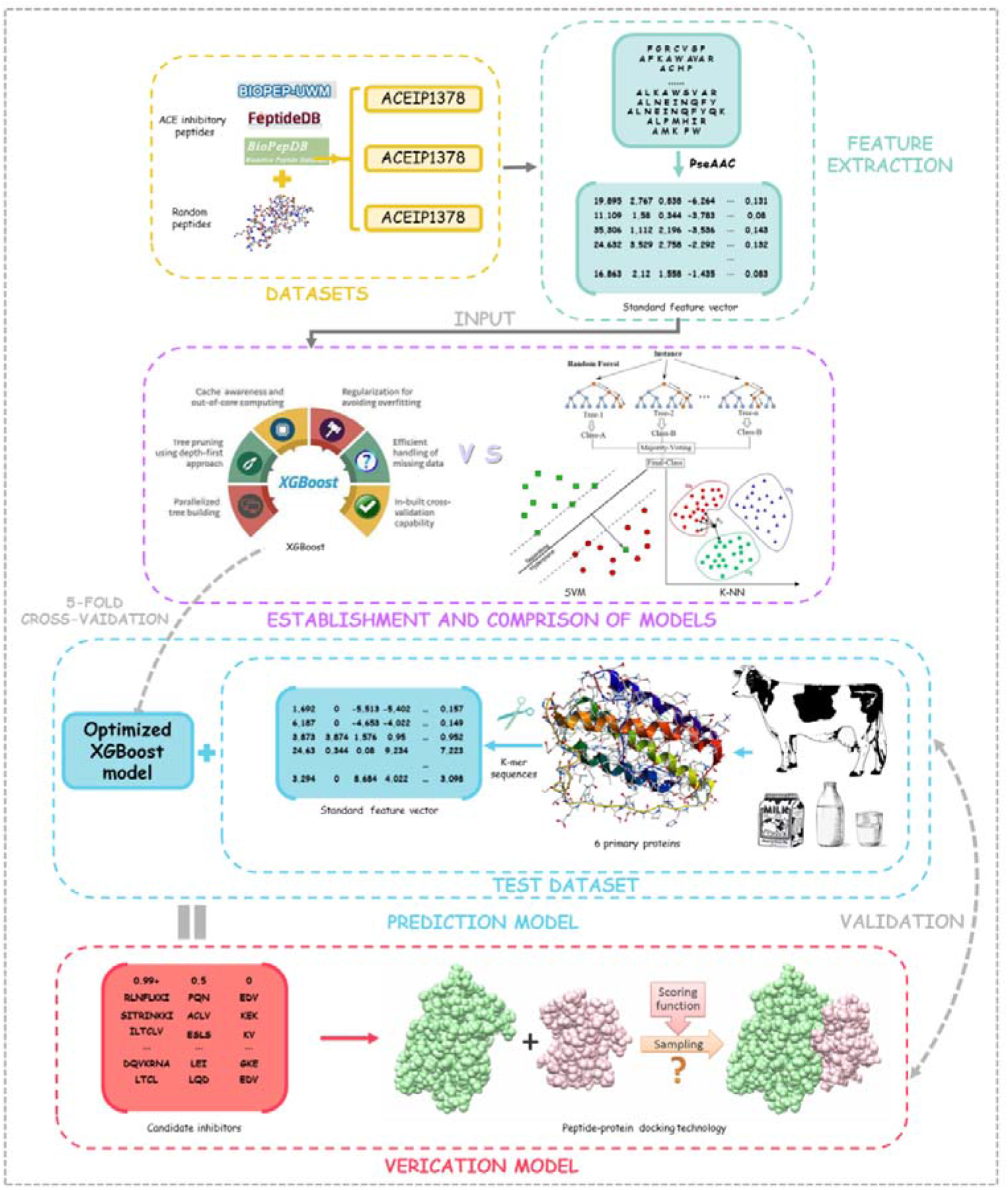
The flow chat of data analysis

## 2 Method and Materials

### 2.1 Training Data and Test Data

In this study, the database of ACE inhibitory peptides for training were obtained from three public databases (BIOPEP-UWM [22], FeptideDB [23–34], and BioPepDB [35]). After screening, 107, 689 and 1653 peptide sequences which were unambiguously tagged as antihypertensive peptides were selected as the positive samples. According to the same strategy employed in previous studies [36,37], we randomly selected the peptides from the UniProt library as the negative samples, which met the equation 1–2 and had the similar average sequence length with the positive samples. The positive samples from there public databases and their corresponding negative samples composed our new datasets, which were named as ACEIP214, ACEIP1378 and ACEIP3306, respectively.

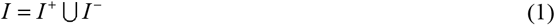

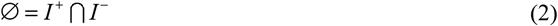

*I*^+^represented the positive samples of antihypertensive peptide, *I*^−^ represented the negative samples, and *I* represented the whole dataset. There was no overlap between *I*^+^ and *I*^−^.

Furthermore, with the aim to test the prediction ability of the model in real situation, 6 key proteins in bovine milk were selected from UniProt protein library, their specific information is shown in **Table 1**. According to the length of peptides in the training datasets, the lengths of created peptide sequence with < 10 were selected to test. Using cycle cutting algorithm, totally more than 10000 k-mer peptides (k = 2, 3, …, 9) [38] were generated from the 6 proteins as the test dataset.

**Table 1:**
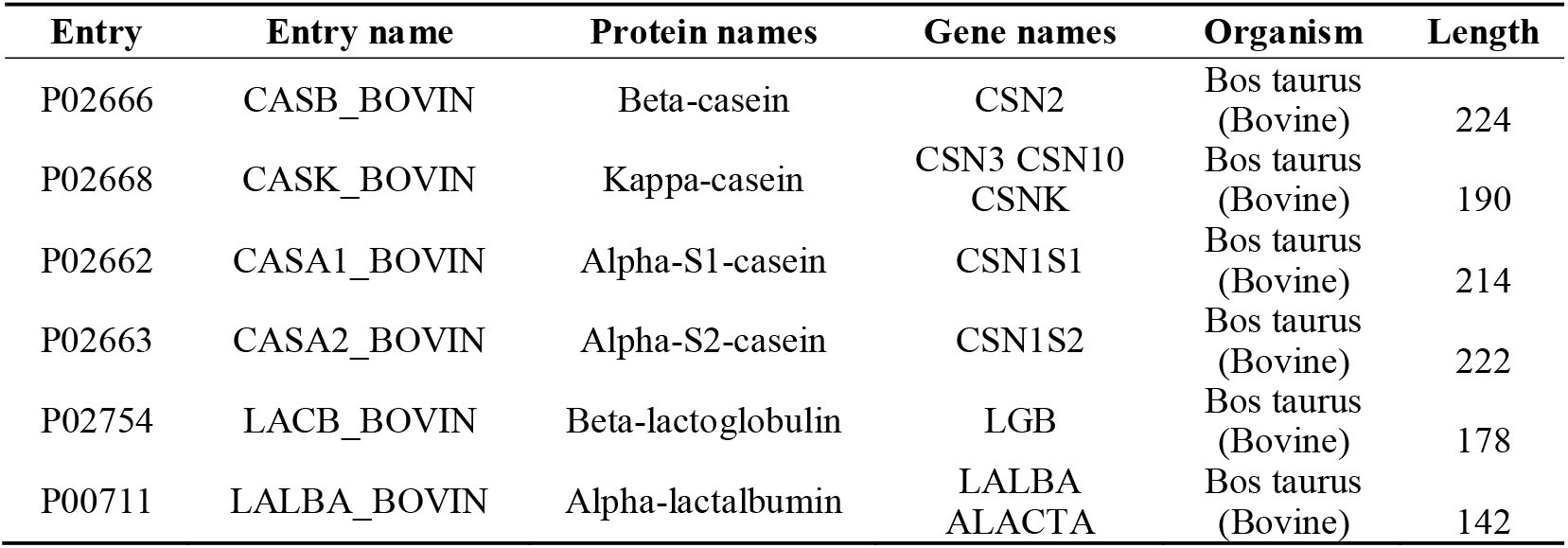
The information of the six key proteins in bovine milk obtained from UniProt.

| Entry | Entry name | Protein names | Gene names | Organism | Length |
| --- | --- | --- | --- | --- | --- |
| P02666 | CASB_BOVIN | Beta-casein | CSN2 | Bos taurus (Bovine) | 224 |
| P02668 | CASK_BOVIN | Kappa-casein | CSN3 CSN10 CSNK | Bos taurus (Bovine) | 190 |
| P02662 | CASA1_BOVIN | Alpha-S1-casein | CSN1S1 | Bos taurus (Bovine) | 214 |
| P02663 | CASA2_BOVIN | Alpha-S2-casein | CSN1S2 | Bos taurus (Bovine) | 222 |
| P02754 | LACB_BOVIN | Beta-lactoglobulin | LGB | Bos taurus (Bovine) | 178 |
| P00711 | LALBA_BOVIN | Alpha-lactalbumin | LALBA ALACTA | Bos taurus (Bovine) | 142 |

### 2.2 Representation of Peptide Sequence Feature

As is widely recognized, the functions of protein largely depend on the 3D structure and some key residues called the reaction center. The 3D structure and the key residues of protein are fundamentally based on the amino acids sequence, indicating that it is possible to infer the function of peptide from amino acids sequence. Additionally, the research of extracting features and representing peptides attract wide interests since the rise of machine leaning technology and several methods has been put up such as AAC, PseAAC (modified algorithm based on classic AAC) and binary profile of patterns (BPP). In present study, PseAAC was adopted as the feature extraction method of peptides.

When encoding the peptide sequence with PseAAC, each peptide can be represented by a vector with 20+iλ dimensions, where i denotes the number of properties of the amino acid taken into consideration and λ is a coefficient that determines the distance of the interacted amino acids (if λ equals 1, only the interaction between adjacent amino acids is considered). Thus, PseAAC is a kind of comprehensive encoding method that includes information of both internal composition and external interaction of the amino acids. The type II PseAAC was applied in our study. In our study, properties of amino acids we chose were hydrophobicity, hydrophilicity, mass, pK1 (α-CO2H), pK2 (NH3) and pI (at 25 □). When *λ* was set to 1, the weight factor ω was set to 0.05.

### 2.3 Machine Learning Algorithms

#### 2.3.1 Extreme Gradient Boosting

XGBoost belongs to one of the boosting algorithms, the idea of which is to integrate many weak classifiers together to form a strong classifier. As a boosting tree model, XGBoost is a powerful classifier composed of many single tree models. The sum of the prediction values generated by each individual in the K-tree (K is the number of single trees) was used in XGBoost model. That is to say, the scores of the corresponding leaf nodes of each tree are pooled as the prediction value of one sample in the XGBoost system. Furthermore, during each iteration of prediction, a new function will be introduced to minimize the objective function as much as possible. Except for linear classifiers employed in XGBoost model, a regular term to the cost function is also added to control the complexity of the model. Moreover, in the contrary with other ensemble learning methods which only use the information of the first-order derivative, the first- and second-order derivatives are considered together and the second-order Taylor expansion of the cost function is executed in XGBoost. XGBoost is a more complicated model than random forest (RF) and compatible with column subsampling, thus making it outperform RF model on training loss, despite of being subject to over-fitting likewise. It is worth noting that XGBoost model has been widely used in various fields due to the superiority of its principle [39,40]. In our study, the PseAAC algorithm was used to extract the primary structural, physical and chemical features of the samples, then the benchmark feature vectors were input into the XGBoost model to train, and thereafter the prediction model was obtained through 5-fold cross-validation.

#### 2.3.2 Support Vector Machine

The basic principle of the SVM model is to find the best separation hyperplane in the feature space, so that the intervals between the positive and negative sample from the training dataset can be maximized. SVM is a kind of classic machine learning algorithm, which belongs to the supervised learning algorithm to solve the two or multi-classification problem. Furthermore, the SVM has been employed to solve nonlinear problems with the introduction of kernel function. Currently, the SVM model has been utilized for peptide prediction as well. For example, Sharma et al. used SVM to predict tumor homing peptides and achieve a maximum prediction accuracy of 86.56% [37]. Yi et al. also adopted SVM as one of the machine learning models for predicting anti-cancer peptides (ACP), resulting in a maximum prediction accuracy of 77.50% [16]. In present study, the parameter-optimized SVM model was also employed to predict ACE inhibitory peptides based on the extracted features of peptide sequence.

#### 2.3.3 Random Forest

RF is a typical algorithm of Bagging-type ensemble learning, which integrates multiple weak classifiers to improve the overall accuracy and generalization ability of the whole model. Though the RF model has been adopted in several research to recognize bioactive peptides [41,42], it undermines when dealing with the unbalanced data of the peptides with high-dimension characteristic. In present study, RF model was also employed to compare with the other algorithms.

#### 2.3.4 K-Nearest Neighbor (K-NN)

As one of the classic algorithms in machine learning, K-NN classifier is widely adopted in various research topics because of its relatively simple principle. It calculates the distance between the new data and the training data, and then selects k (k ≥ 1) closest neighbors to make classification or regression. The K-NN model has been applied in protein recognition [43]. It is convenient to use the K-NN model due to its simple theory and relatively lower complex training process. However, it is inevitable to face the low interpretability and prediction accuracy of rare categories when the sample is unbalanced. In present study, the K-NN algorithm after parameter optimization was also employed to compare with the other machine learning methods in the task of ACE inhibitors screening.

### 2.4 Performance Evaluation of Models

The 5-fold cross-validation was executed in our study, and the results were displayed by the commonly used evaluation criteria, including Accuracy (Acc), Sensitivity (Sens), Specificity (Spec) and Precision (Prec) (The specific definition of these indicators were showed in equation 3–6) and the area under the receiver operating characteristic curve (AUC).

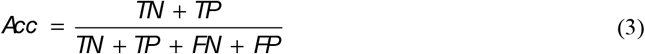

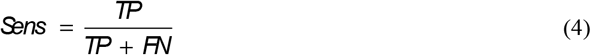

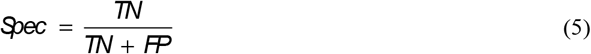

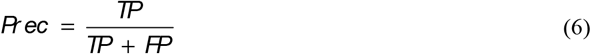

Where TN represents the true negative number, TP signifies the true positive number, FN denotes the false negative number, and FP stands the false negative number.

### 2.5 Prediction Model and Peptide-Protein Docking Verification

To test the prediction ability of our ACE inhibitory peptide model in the real situation, the optimal model obtained from training process was utilized to do high-throughput and rapid peptide screening of our test dataset (comprised over 10000 short peptides cutting from the key proteins rich in bovine milk). The test experiments were performed parallelly for three times, and possibility of positive peptide was calculated. When the possibility of one peptide is over than 99.00% for all of the three times, the peptide can be recognized as the one with anti-hypertensive activity in our study. Furthermore, in order to discover the difference between positive and negative peptide predicted in present study, two groups of peptides with possibility of 0.00% and 50.00% respectively were selected as the control groups.

The screening results of our model were further verified via peptide-protein docking technology. With help of virtual screening technology, discovering new inhibitors is becoming a common practice in modern drug discovery [44]. Furthermore, structure-based virtual screening approach is widely employed in this field due to its cost-effective and time-saving advantages. In our research, virtual screening was applied to validating the prediction results of our models. HPEPDOCK Server was selected to carry out our virtual screening task due to its outstanding performance and accurate result [45–54]. The targets of the verification experiment were the 3 group peptides with differentiable possibility of anti-hypertensive peptide (the verified dataset was obtained from the above part of prediction). Considering the fact that the reaction center of ACE is clearly known, it is reasonable to judge the docking result by the docked free energy (measured as the docking scores by this server). Theoretically, peptides that are fixed to the pocket of reaction center with lower affinity energy are more likely to be the inhibitors and vice versa.

## 3 Results

The computer was configured as CPU Intel Core I7-6700HQ, 3.5 GHz, 4 GB of memory and the experimental programming was implemented in Python 3.8. The composition of all 20 amino acids in peptides were counted and compared in our datasets, and then the benchmark feature vectors obtained via PseAAC algorithm were input into the models. Through cross-validation, the performance of the XGBoost algorithm and some other algorithms were compared, and the optimal model was selected out for subsequent testing. During the testing process, the k-mer algorithm was utilized to cut the primary six proteins in the bovine milk, and short peptides with different lengths were used as the test dataset, and then the probability of ACE inhibitory ability obtained by our optimized model were generated as the prediction results. In order to show the reliability of our method, peptide-protein docking technology was exploited to verify the prediction results.

### 3.1 Distribution of Amino Acids in the Datasets

We counted and compared the composition and amino acids distribution of positive samples, negative samples and all samples in our three benchmark datasets, respectively (Shown in **Figure 2**). It is obvious that Pro and Leu appeared frequently in ACE inhibitory peptides, while Cys, Met and Trp were rare in these peptides. Thus, Pro and Leu were inferred to make a difference in inhibiting the ACE activity.

**Figure 2:**
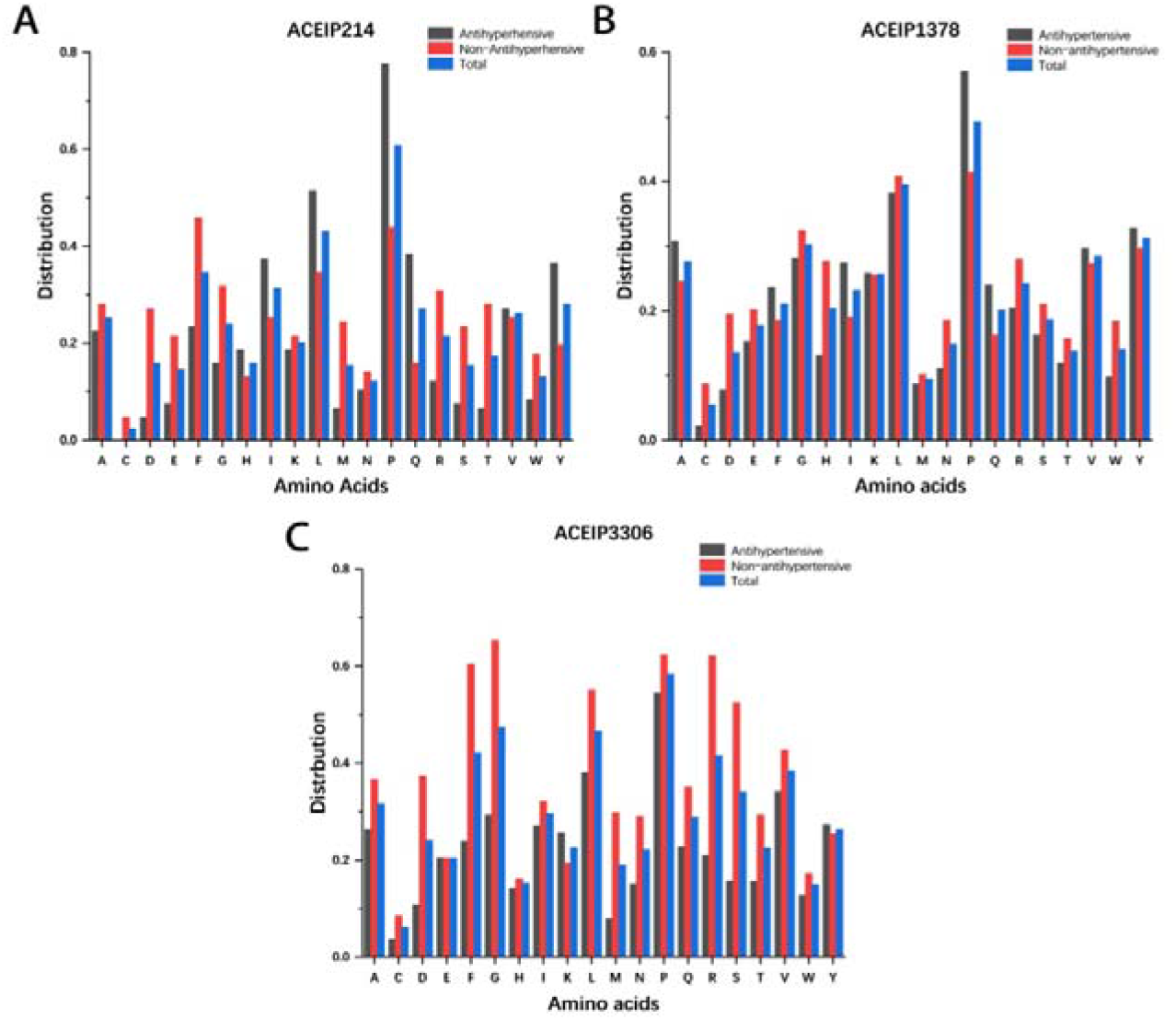
The frequency distribution of various amino acids in peptides from the three datasets ACEIP214 (A), ACEIP1378 (B) and ACEIP 3306 (C)

### 3.2 Cross-Validation Results of XGBoost Model

In present study, the XGBoost model was adopted to execute 5-fold cross-validation based on the three datasets ACEIP214, ACEIP1378, and ACEIP3306 and the results are showed in **Table 2**. The best performance of XGBoost model was achieved in ACEIP3306, with the mean accuracy of 0.9841, the average sensitivity of 0.9831, the average specificity of 0.9806, and the average accuracy of 0.9850, which reflected the excellent performance and strong generalization ability of XGBoost algorithm. The values of corresponding indicators on ACEIP 214 and ACEIP 1378 were deviated from ACEIP3306 a little bit and the underlying reasons are listed as following: firstly, the size of the dataset will affect the accuracy of the model. In general, the huge data enables the model to excavate and grasp the deep features, making the prediction more rational, so that the model can achieve better results with the increase of the size of the dataset; Secondly, the intrinsic discrepancies between the three datasets may impact the performance of the model. It is noted that our datasets came from different database libraries. In fact, there exit slight discrepancies between different database libraries, especially for their measures or standards to judge whether a peptide is antihypertensive or not. Although such discrepancies may be very minute, they do attribute to the deviation on the results.

**Table 2:** Performance of XGBoost model on different datasets.

| Name of Dataset | Acc | Sens | Spec | Prec |
| --- | --- | --- | --- | --- |
| ACEIP214 | $0.9298 \pm 0.0134$ | $0.9310 \pm 0.0159$ | $0.9286 \pm 0.0201$ | $0.9288 \pm 0.0086$ |
| ACEIP1378 | $0.9328 \pm 0.0156$ | $0.9229 \pm 0.0221$ | $0.9430 \pm 0.0107$ | $0.9418 \pm 0.0093$ |
| ACEIP3306 | $0.9841 \pm 0.0052$ | $0.9831 \pm 0.0028$ | $0.9806 \pm 0.0044$ | $0.9850 \pm 0.0049$ |

Notes: the above results show the average value of each indicator and the corresponding 95% confidence interval. Additionally, Acc, Sens, Spec and Prec represent accuracy, sensitivity, specificity and precision, respectively.

In order to comprehensively display the performance of the model, the receiver operating characteristic curve (ROC) and AUC were introduced as significant indicators. The corresponding curves of XGBoost model are exhibited in **Figure 3**. It was obvious that significant differences existed among the AUC value of different datasets. Among them, ACEIP3306 had the largest AUC of 0.9428, followed by the 0.9277 of ACEIP1378 and the 0.8248 of ACEIP214.

**Figure 3:**
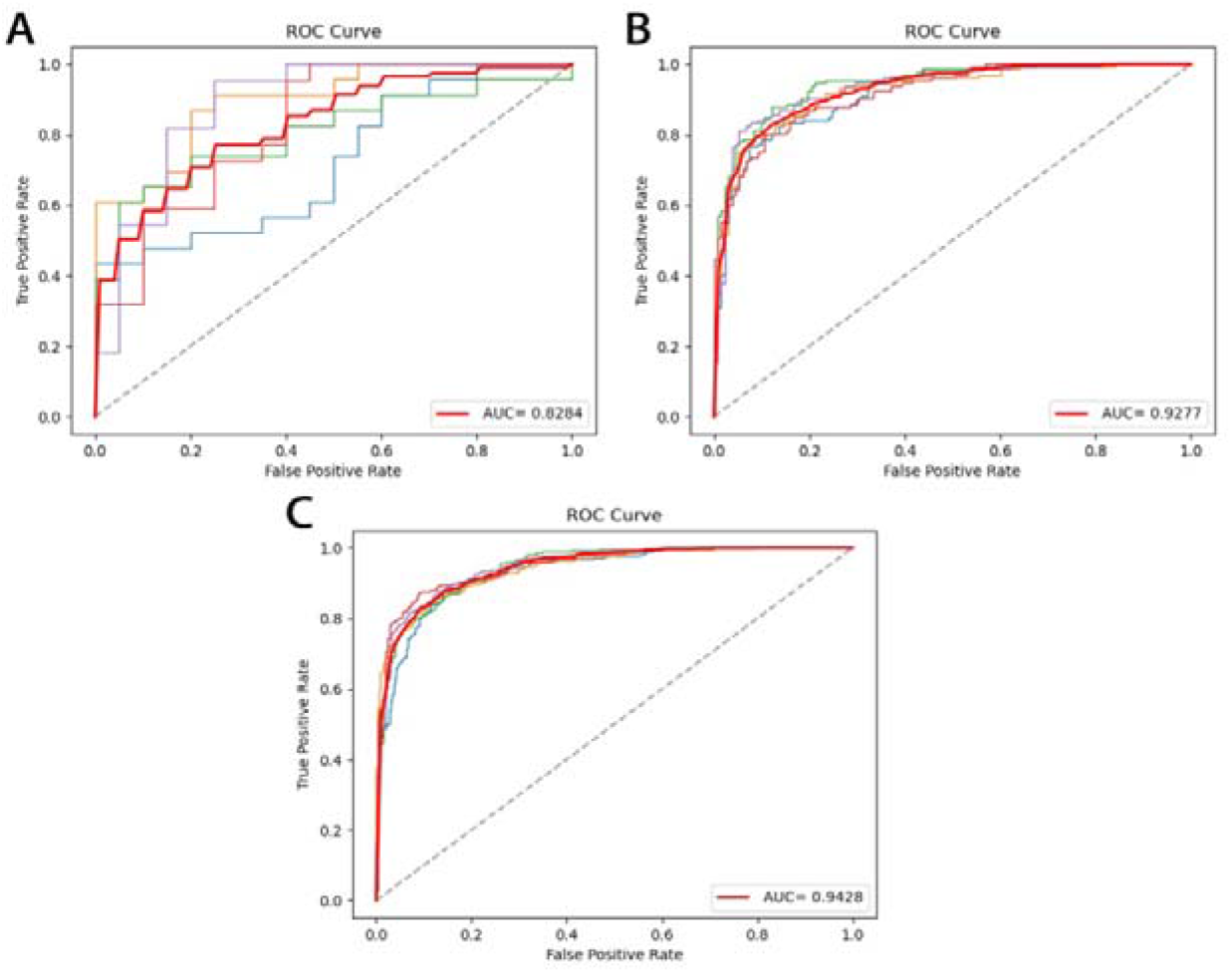
Performance of XGBoost model on the datasets ACEIP214 (A), ACEIP1378 (B) and ACEIP3306 (C)

### 3.3 Cross-Validation Results of Other Models

We further constructed the models with other machine learning methods and obtained the results of corresponding indicators (displayed as the 95% confidence interval of the average) through 5-fold cross-validation (shown in **Figure 4**). Our results showed that the XGBoost model exhibited the remarkable ability when compared with the classic machine learning algorithms. Furthermore, we also found that the performance of RF and K-NN models were better than SVM, and their mean accuracy of ACEIP3306 was higher than the other two datasets, which were up to 0.887 and 0.897. RF is a classic Bagging ensemble model which has a strong generalization ability regardless of the data scale, and the parameter n_tree was set to 20 in our study. Using simple principle and several parameters, K-NN algorithm can reduce the complexity and difficulty to establish and optimize the model, and the parameter k was set to 4 in the present study. Though the accuracy of SVM based on ACEIP3306 was significantly lower than RF and K-NN, its performances based on ACEIP1378 and ACEIP214 were similar with RF and K-NN, indicating that SVM is more suitable for training dataset with a small scale. Usually, the quadratic programming by speculation is used in SVM to solve the support vector, which is time and space-consuming because solving quadratic programming is involved with the calculation of the mth order matrix (m is the number of samples). When m is large, lots of machine memory and computing time are needed by the storage and calculation of the matrix

**Figure 4:**
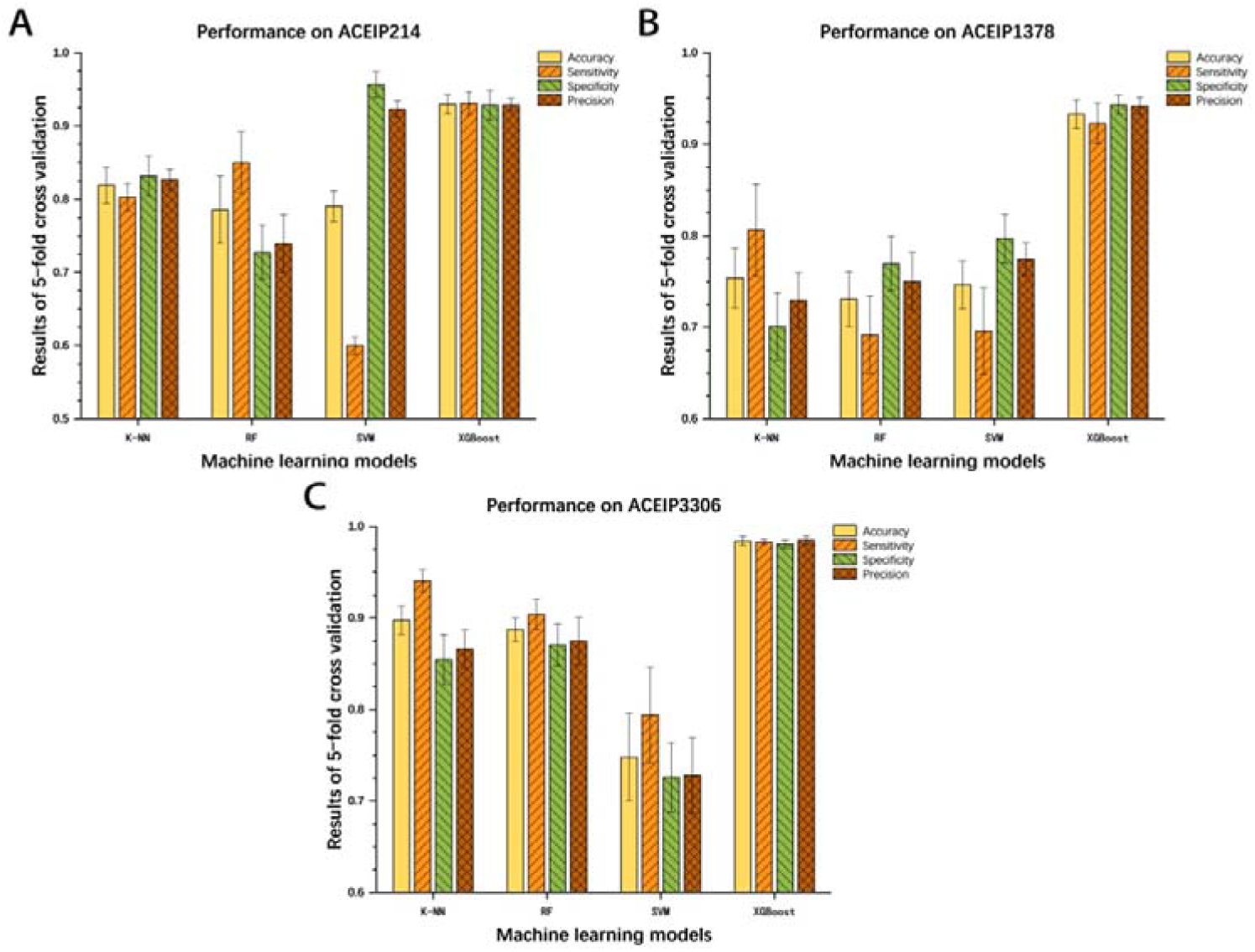
Comparisons and evaluations of different machine learning models on datasets ACEIP214 (A), ACEIP1378 (B) and ACEIP3306 (C)

The ROC curves and AUC values of the other three classic machine learning methods on different datasets are showed in **Figure 5**. Comparing with the performance of XGBoost model, all three of classic machine learning algorithms had significant lower values of AUC based on the three datasets, which supporting outstanding generalization and superior ability of XGBoost for ACE inhibitory peptides screening task. In addition, the AUCs of RF (0.6967, 0.7674 and 0.9373) were higher than SVM (0.5966, 0.6768 and 0.8439) and K-NN (0.6846, 0.7381 and 0.9003) on the same datasets, respectively.

**Figure 5:**
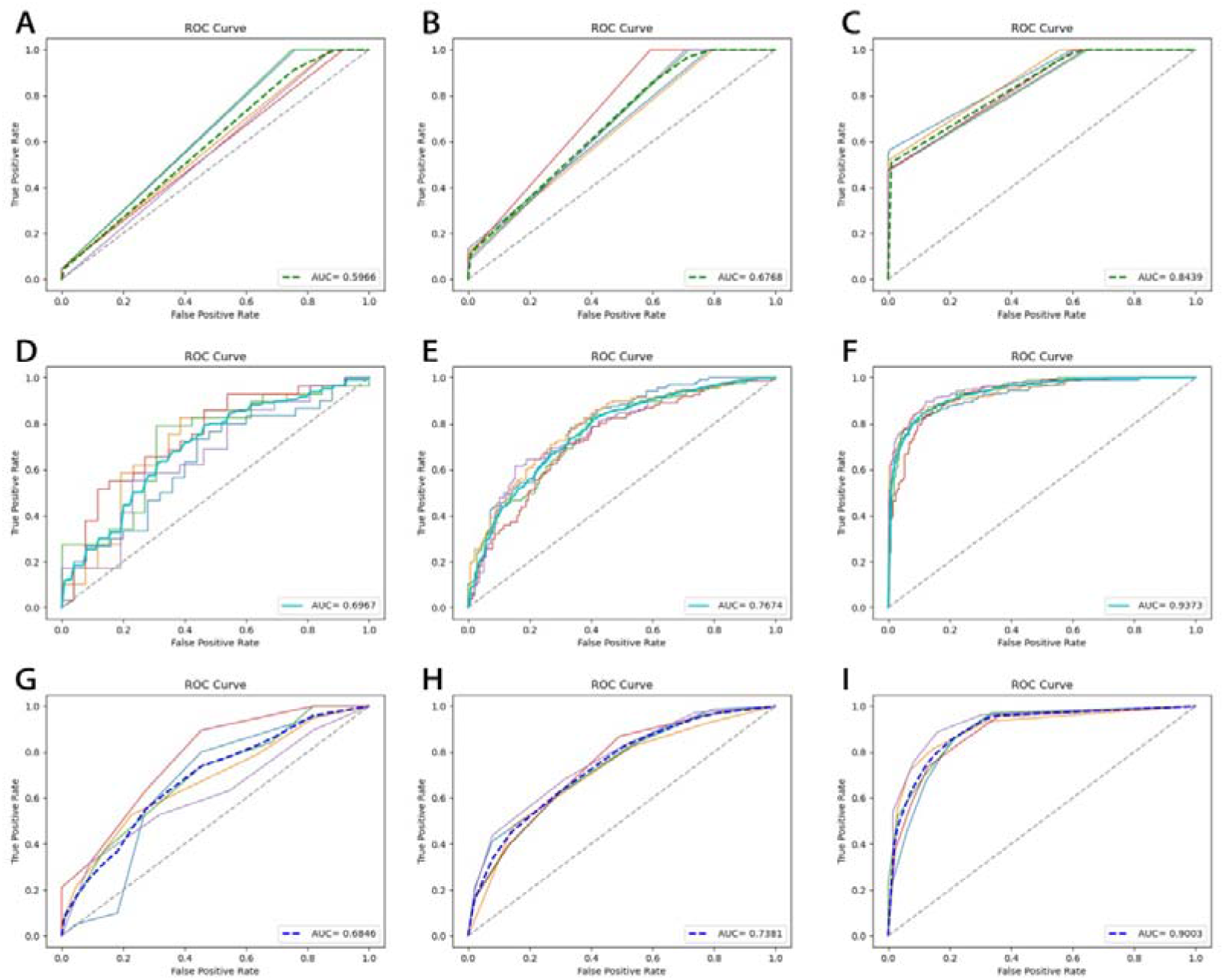
The performance of machine algorithms on different datasets. The performance of SVM model on ACEIP214 (A), ACEIP1378 (B) and ACEIP3306 (C), The performance of RF model on ACEIP214 (D), ACEIP1378 (E) and ACEIP3306 (F), the performance of K-NN model on ACEIP214 (G), ACEIP1378 (H) and ACEIP3306 (I)

### 3.4 Prediction Model and Protein-peptides docking Verification

In present study, over 10000 k-mer (k=2, 3, …, 9) peptides were obtained by cutting the 6 key proteins in bovine milk. Based on the performance of the models, we further chose the XGBoost model (obtained by training ACEIP3306) to predict ACE inhibitory peptides. After importing into the model, the results of three parallel experiments and the prediction time was recorded. In the end, three groups of candidate inhibitory peptides were generated based on their probability value. In present study, the group with probability values of prediction over 0.9900 for 3 times was recognized as candidate inhibitory peptides, and the other two groups with 0.00 and 0.5000 were chosen as the controls (Details are exhibited in **Table 3**). Based on the existing computing equipment, the screening process consumed 903.4 s in total (average 0.082 s/peptide). In order to verify the efficiency and feasibility of our prediction model, the peptide-protein docking was conducted on the candidate ACE inhibitory peptides. The docking results (represented as relative free energy by the platform) and some visual 3D structural diagrams were displayed in **Table 3** and **Figure 6**. There was a significant difference of the relative free energy between the candidate inhibitory peptides and the controls in other two groups (P < 0.05), indicating that the affinity between the candidate inhibitory peptides and ACE enzyme was evidently greater. It is worth noting that the speed of the working platform was largely affected by the length of the peptide participating in the docking task, and its docking speed is up to 480.0~1680.0 s/peptide. Therefore, our method is able to achieve high-accuracy, high-throughput, and high-speed screening tasks by comparison with the traditional computer drug-assisted screening technology.

**Table 3:**
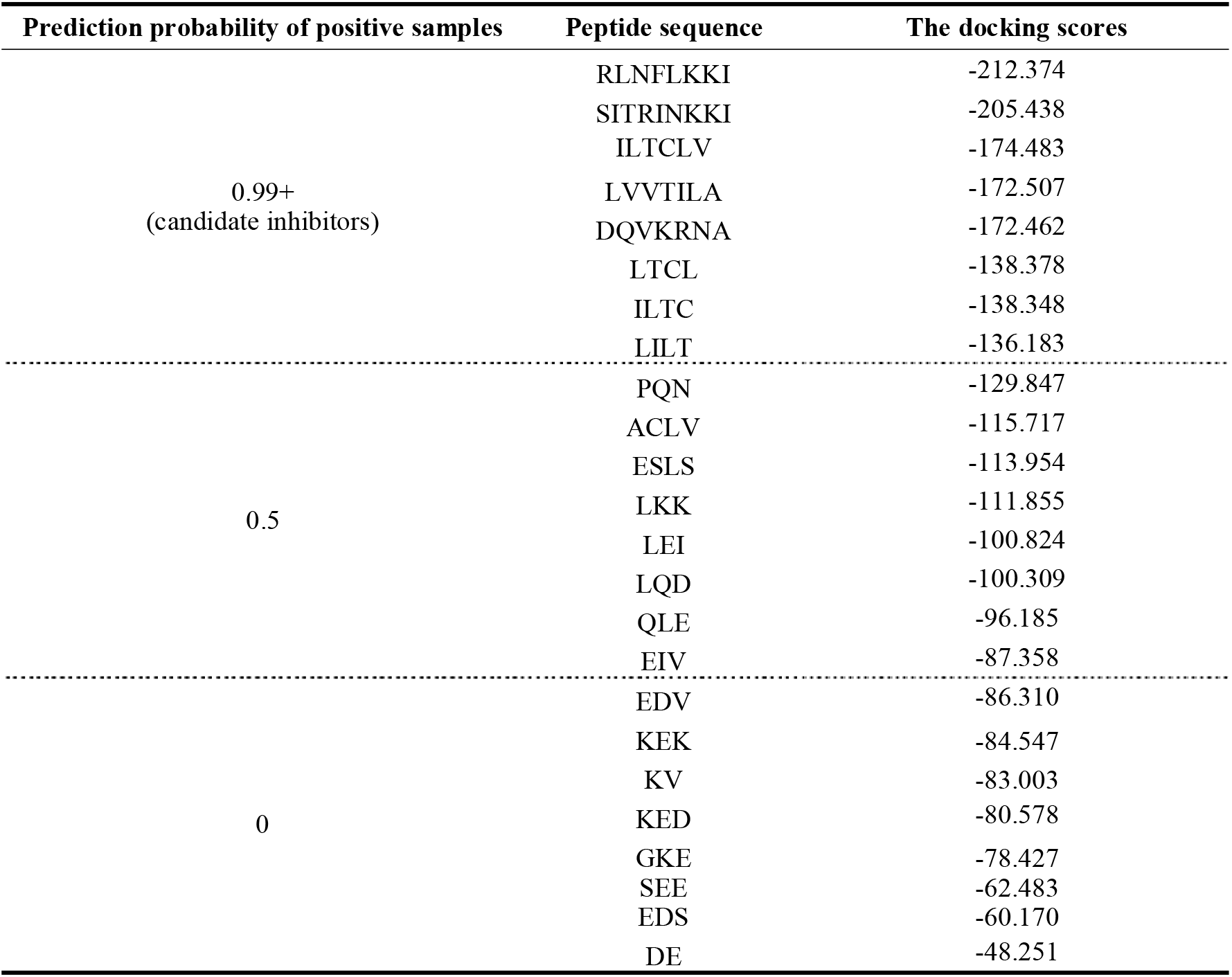
Results of screening and the corresponding docking scores (relative free energy).

**Figure 6:**
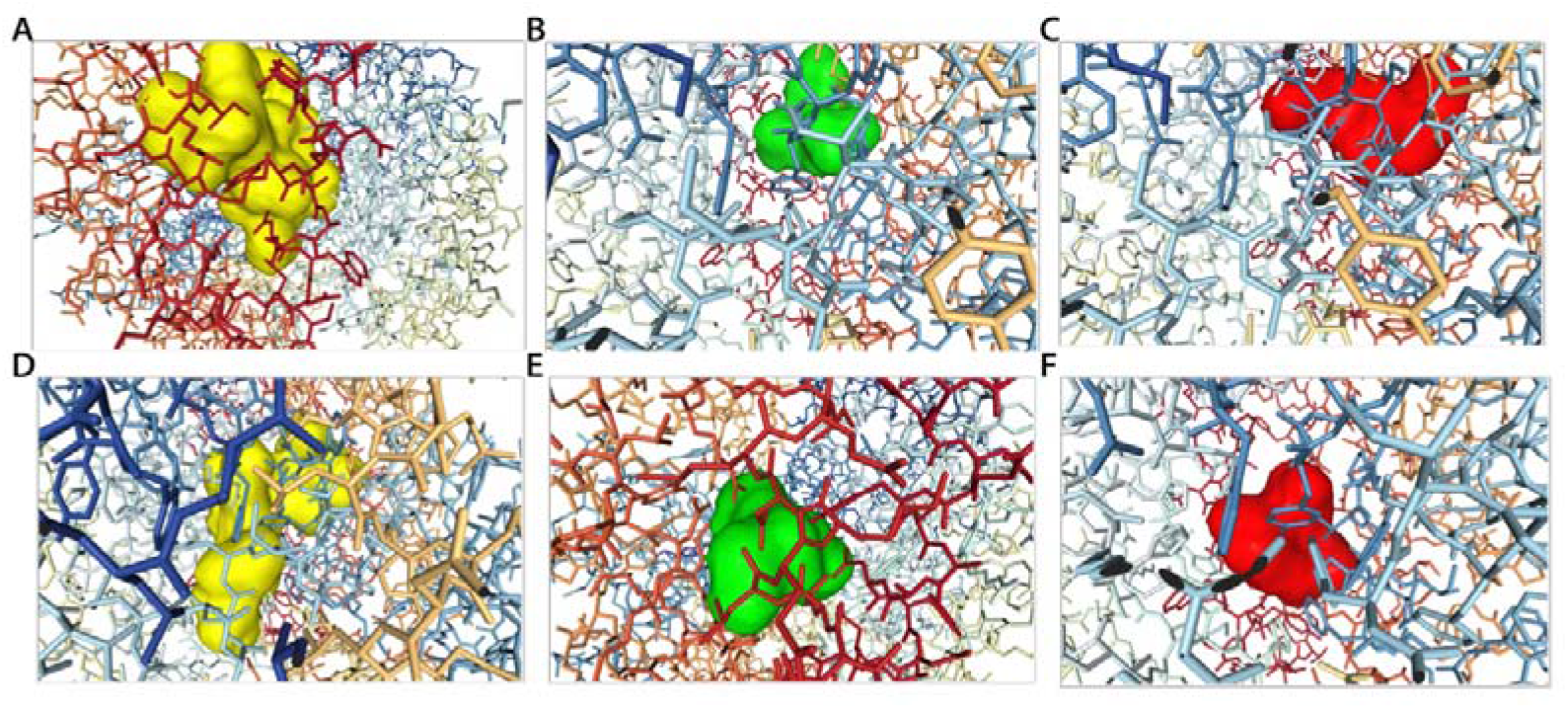
The 3D structural diagrams of protein-peptide docking of RLNFLKKI (A), PQN (B), EDV (C), SITRNKKI (D), ACLV (E), KEK (F)

## 4 Discussion

In this study, to screen ACE inhibitory peptides with high throughput, precision and speed, 4 machine learning models were established and compared based on three benchmark datasets and the XGBoost model was selected as the one with best performance. Furthermore, to verify the reliability and utility of the model in the real situation, a test dataset with over 10000 short peptides cutting from the 6 key proteins rich in bovine milk was structured and the optimal model was adopted to predict the ACE inhibitory. Meanwhile, the peptide-protein technology was employed to verify the validation of the results of prediction. Our study validated the superiority of XGBoost model in the task of ACE inhibitory peptides. In the other aspect, the new idea that using machine learning method to explore and discover antihypertensive peptides from food protein with known structure is proved to have potential value.

Three benchmark datasets (ACEIP214, ACEIP1378, ACEIP3306) with the same number of positive ACE inhibitory peptide samples and negative ones were established from the selected three public databases. The results indicated that ACEIP3306 performed the best in 5-fold cross-validation process regardless of the algorithms we chose, followed by ACEIP1378, and finally ACEIP214. It was speculated that the size scale of the training dataset has affected the learning ability of the corresponding algorithm. That is to say, increasing the size of the dataset may enable the model to exert the deeper features, thus improving its prediction accuracy [55]. Limited by the current reports of ACE inhibitory peptides, we can only do the training and prediction in a small range. In the future, the database should be expanded to realize the goal of ACE inhibitory peptides screening in a larger range. Additionally, slight differences between different databases have existed in the rule and standard to justify whether a given peptide is antihypertensive or not, which may attribute to the difference of cognitive ability of the models between the datasets. As a result, unifying and perfecting the specific standards and protocols may be beneficial to improve the generalization of the model.

As a novel Bagging ensemble learning algorithm, it is innovative to adopt XGBoost in the task of antihypertensive peptides, for despite the algorithm has been proven to have excellent performance in other fields [56,57]. In our study, the primary structural feature of the peptides represented by the PseAAC was inputted to the XGBoost model. Using the 5-fold cross-validation, Acc, Sens, Spec, Prec and AUC were calculated as the indicators to evaluate the models. Meanwhile, three classic machine learning methods (SVM, RF, and K-NN) were employed to screen and predict properties of peptides. The results showed that the XGBoost model performed better than the other algorithms based on all of the three datasets. More specifically, the regularization terms are introduced into the XGBoost model to control the complexity of the model, which not only simplifies the learning model but also prevents overfitting. On the other hand, XGBoost draws the idea of RF that supports column sampling, which reduces the overfitting and calculation [58]. Besides, for the samples that occasionally miss part of the features, XGBoost can automatically learn its splitting direction. The XGBoost model also uses a greedy algorithm to enumerate all possible split points, which benefits the generating of the optimal tree structure.

The test dataset comprised of over 10000 sequences from 6 key proteins in bovine milk were used to verify the reliability of our optimal model (training with dataset ACEIP3306). A possibility value represented as the ACE inhibitory degree was obtained for every sequence thorough the XGBoost model. The results of three parallel experiments were consistent with the peptide-protein docking results, thus proving the feasibility of machine learning method as a novel auxiliary tool for ACE inhibitory peptide screening. It is worth noting that the screening speed of our method was remarkable faster than the traditional docking technology, indicating its potential to achieve high-throughput, rapid and accurate screening task. It should be emphasized that 6 key proteins in bovine milk was used as the raw material, and some short peptides cleaved from them were predicted to have antihypertensive properties by our model. The strategy give us enlightenment of employing machine learning algorithms to rapidly screen and obtain peptides with specific functions from natural foods in future.

Furthermore, compare this work with previous related reports. In the research of ACE inhibitors screening, Ya et al. [12] used SVM algorithm and ligand-based QSAR model to predict ACE inhibitors, resulting in an accuracy of 0.8874. Guan et al. [13] successfully established the QSAR model using orthogonal signal correction combined with SVM (OSC-SVM) through 268 peptides, which showed relatively excellent fitting accuracy and generalization ability in the task of predicting ACE inhibitory peptides. In the screening tasks of other targets, for example, Cai et al. [14] employed machine learning models such as Plain Bayes and recursive partition (RP) algorithms to predict DPP-IV inhibitors; they established 247 sub-models based on 1307 known DPP-IV inhibitors, and the final overall prediction accuracy exceeded 80%. Chandra et al. [15] also designed an SVM algorithm to predict DPP-IV inhibitors with the Matthew correlation coefficient in the external test set of 0.883, and they have further applied the method to Web programs. Yi et al. [16] screened the ACP utilizing long short-term memory (LSTM), which achieved better performance than traditional machine learning method. However, in contrast, our research has achieved better predictive capabilities due to the use of superior algorithm and larger datasets.

It should be acknowledged that flaws still existed in our algorithm. Due to the lack of definitely validated non-ACE inhibitory peptides, the negative samples in our datasets were replaced by random peptides, which inevitably mixed some ACE inhibitors. This possibility will doubtlessly reduce the prediction accuracy of the model. In addition, the k-mer sequences in our test datasets gained from proteins in bovine milk were obtained by theoretical segmentation algorithms without considering their biological activity and feasibility in real situation. As a result, the feasibility of extracted antihypertensive peptides from natural food still depends on the development and advance in enzyme cleavage technology in the near future.

## 5 Conclusions

In this study, a method of utilizing PseAAC to extract primary structural features of peptides and then establishing XGBoost model to predict their antihypertensive properties was proposed. This method achieved excellent performance in the task of antihypertensive peptides screening with high throughput, great precision and rapid speed. In order to fully present the superiority and robustness of the method in processing different datasets, three baseline machine learning methods (SVM, RF and K-NN) were introduced for comparison on three datasets with different size scales. In our experiment, the XGBoost model demonstrated its superiority through 5-fold cross-validation with the highest mean accuracy of 0.9841 and the highest average AUC of 0.9428. In addition, the introduced peptide-protein docking technology has verified the reliability and efficiency of our model compared with the traditional strategies, its properties of rapid speed and high throughput also provided us with promising prospect that many ACE inhibitory peptides can be extracted from natural food with skillful and time-efficient method in the near future. Moreover, the study offered us with a more reliable auxiliary method for the in vitro and in vivo experiments in future.

## Author Contributions

Conceptualization, L.W. and D.N.; methodology, L.W.; software, L.W.; validation, D.N. and X.W.; formal analysis, D.N.; data curation, X.W.; writing—original draft preparation, L.W., Y.X., and Q.S.; writing—review and editing, L.W.; visualization, X.W. and Q.S.; project administration, Y.X.; funding acquisition, Y.X. All authors have read and agreed to the published version of the manuscript.

